# Efflux Pumps Represent Possible Evolutionary Convergence onto the Beta Barrel Fold

**DOI:** 10.1101/268029

**Authors:** Meghan Whitney Franklin, Sergey Nepomnyachiy, Ryan Feehan, Nir Ben-Tal, Rachel Kolodny, Joanna S.G. Slusky

**Author notes:** Corresponding author and lead contact: Joanna S.G. Slusky.

## Abstract

There are around 100 types of integral outer membrane proteins in each Gram negative bacteria. All of these proteins have the same fold—an up-down β-barrel. It has been suggested that all membrane β-barrels other than lysins are homologous. Here we suggest that β-barrels of efflux pumps have converged on this fold as well. By grouping structurally-solved outer membrane β-barrels (OMBBs) by sequence we find evidence that the membrane environment may have led to convergent evolution of the barrel fold. Specifically, the lack of sequence linkage to other barrels coupled with distinctive structural differences, such as differences in strand tilt and barrel radius, suggest that efflux pumps have evolutionarily converged on the barrel. Finally, we find a possible ancestor for the OMBB efflux pumps as they are related to periplasmic components of the same pumps.

## Introduction

All bacterial outer membrane proteins save one (Dong et al., 2006),are right-handed, up-down β-barrels with the N and C termini facing the periplasm. This extreme topological homogeneity has given rise to questions of outer membrane β-barrel (OMBB) evolutionary origin. Specifically, did this fold arise from divergent evolution of a single common ancestor or did multiple ancestors converge onto an identical fold required by the biological and physical constraints of the outer membrane? In support of divergent evolution, Remmert et al. propose that the strand number diversity results from amplification of an ancestral hairpin or double hairpin (Remmert et al., 2010).

A useful counter example to the well-established hypothesis of divergence of all OMBBs from a common ancestor is the existence of membrane barrels, such as alpha hemolysin and the leukocidins, which are not localized in the outer membrane of Gram negative bacteria. Unlike the OMBBs, the lysins are exported during cellular warfare to create pores in membranes of other organisms. It has been hypothesized that the lysins have evolutionarily converged to their barrel structure separately from the divergent evolution of the other β-barrels. This hypothesis was based on the distance in sequence, the difference in structure, and on the differences in organisms that produce them (Remmert et al., 2010).

Efforts to document the homology among OMBBs have been frustrated by two factors, 1) the high sequence similarity required for the strands of β-barrels and 2) extreme bacterial sequence variation. First, β-barrel structure and environment collude to enforce common sequence patterns. The β-barrel structure causes half of the positions to be facing the membrane and half of the positions to face the interior which is an (often solvated) pore. Therefore, β-strand sequences organize into a pattern with a polar-nonpolar alternation of residues in each strand (Schulz, 2002). This high level of environmentally enforced sequence similarity means that some barrels may appear sequence similar though they are not homolgous.

Second, β-barrels are also subject to extreme variation through evolution. The Gram negative bacteria is likely about 3 billion years old (Battistuzzi et al., 2004), and bacterial generation can be quick— generally less than an hour, though only 10% of its time is in growth phase. This rapid replication adds another three orders of magnitude, resulting in at least 10^12^ opportunities for introducing genetic variation such as amplification, recombination and accretion of mutations. In addition, membrane proteins are less conserved than soluble proteins, partly because they are more involved in adapting to new environments (Sojo et al., 2016). For outer membrane proteins the lipid-facing side is particularly prone to variation (Jimenez-Morales and Liang, 2011). Ultimately, these factors can result in OMBB proteins diverging beyond sequence recognition which means that the usual rule of thumb of only E-values of less than 10^−3^ (Pearson, 2013) being suggestive of homology may not apply.

To establish homology of OMBBs, previous studies created large databases of sequence similar proteins and then culled those sequences to increase the likelihood of true OMBBs (Remmert et al., 2010), (Reddy and Saier, 2016). Creating these large databases of sequences is extremely useful for teasing out evolutionary relationships within proteins that are related. The benefit of larger databases is that it allows for the creation of a more connected network. However, using these large databases to understand relationships between structure and homology is harder. Without experimental structural determination, structure prediction must be used. Both homology by sequence and predicted structure/topology are reliant on the construction of Hidden Markov Model (HMM) profiles which are sequence dependent. HMMs determine homology using a probabilistic model with the sequences as the inputs. The dependence on sequence based HMMs is especially the case as HMMs are used in all the most successful methods of OMBB structure prediction (Bagos et al., 2005). This leads to a double counting of the sequence relationships that may obscure the very thing we are trying to determine: the relationship between protein fold and protein homology.

Finding a relevant database to determine the homology through combined sequence and structural relationships can be challenging. As structural data for bacterial OMBBs have increased, the annotated databases of protein domains have struggled with whether to place OMBBs into the same or separate groups (Tsirigos et al., 2018). In general, current classification schemes reflect the underlying homologous relationships, but the high degree of sequence divergence makes that difficult to achieve for OMBBs. For example, SCOPe (Fox et al., 2014) artificially defines a class of “membrane and cell surface proteins and peptides”. The OMBBs fall into four different folds in SCOPe, of which the fold “transmembrane β-barrels” is labeled as “not a true fold”. CATH (Sillitoe et al., 2015) strictly classifies OMBBs by overall topology and many of the more recently characterized structures do not yet have a CATH classification. In CATH with the exception of the adhesins like HiaA, the OMBBs fall into the all-β class and then into at least five different architectures. Finally, most of the single-chain OMBBs fall into the same possibly-homologous group of outer membrane meander β-barrels in ECOD (Cheng et al., 2014). However, the multi-chain β-barrels, like the efflux pumps, adhesins, and especially the lysins, are splintered into many different groups.

Here, we studied the evolutionary relationships among OMBB proteins, and present evidence that the outer membrane components of tripartite efflux pump proteins have converged on the OMBB fold. These efflux pump proteins are responsible for expelling metals and most small hydrophobic molecules from bacterial cells. They have recently become a principle focus of scientific interest as they are the primary mode of Gram negative antibiotic resistance. Tripartite efflux pumps expel most classes of antibiotics and biocides including chloramphenicol, tetracycline, macrolides, fluoroquinolones, quaternary ammonium compounds, amino glycosides, triclosan and even the pine oil used in pine scented cleaner (Liu et al., 2010; Poole, 2005). In clinical isolates, expression of tripartite efflux pumps have been found to correlate with resistance (Swick et al., 2011). These pumps extend from the inner membrane, through the periplasm, to the outer membrane. As per their name, tripartite efflux pumps are comprised of three types of proteins, outer membrane factors (OMFs), periplasmic adaptor proteins (PAPs), and inner membrane transporters (IMTs). These efflux pumps utilize the proton motive force of the inner membrane to drive substrate transport (Zgurskaya and Nikaido, 1999).

Due to the fractured membership of the OMBBs in the domain-centric databases, our study relies on creating a bespoke data set to better interrogate the OMBB structural class. By combining structural data with sequence data, we describe results that suggest efflux pump OMBBs may have evolutionarily converged to the OMBB fold. Ultimately, the structural insights described inform our understanding of the mechanism of antibiotic resistance.

## Results

**Figure 1.**
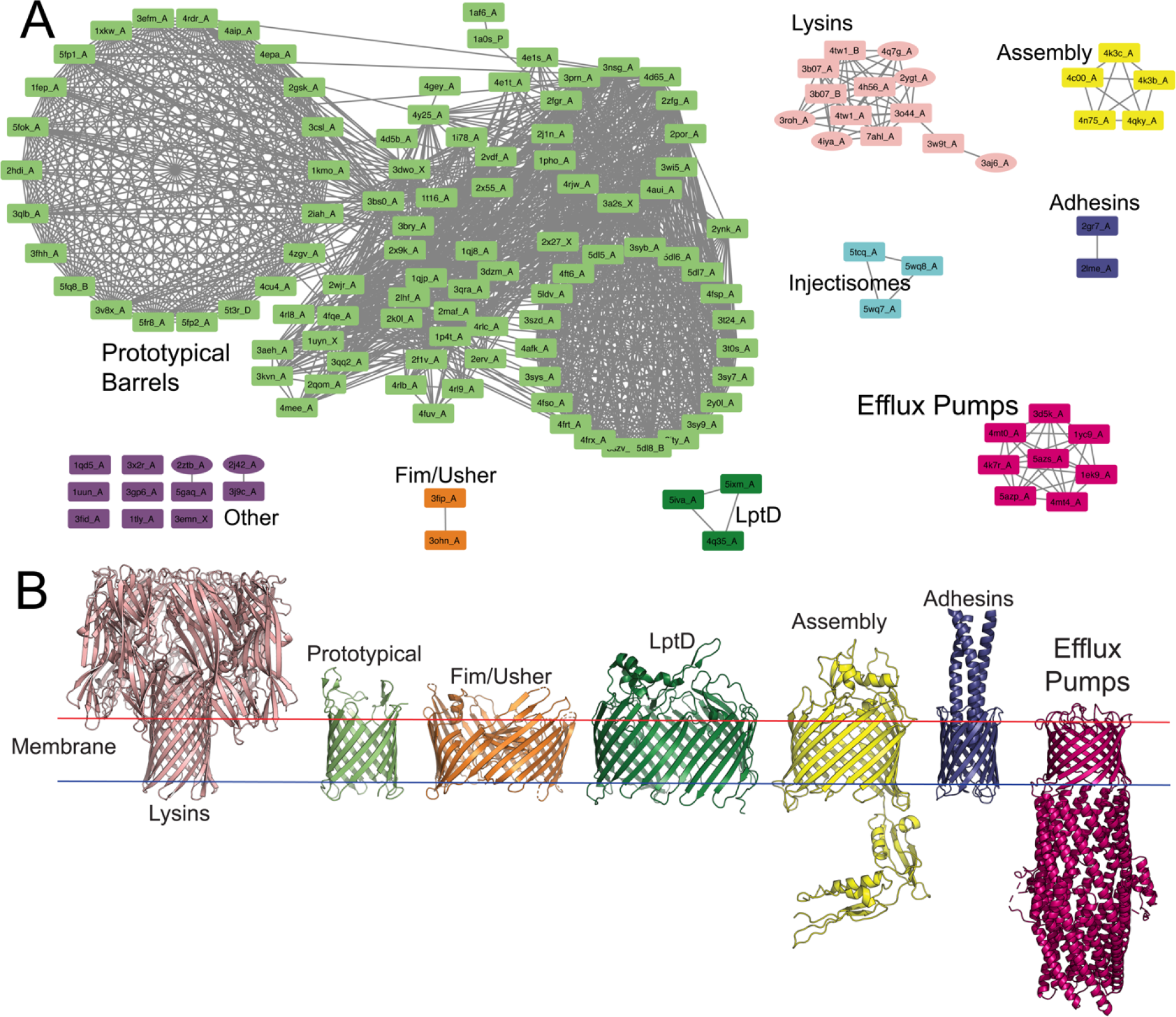
A) Sequence alignments among the membrane beta barrels at E < 1, colored by component; ovals are structures for which only a monomer has been crystalized but the protein is proposed to be a full beta-barrel. The purple group are defined as the barrels for which no alignment to any of the other barrel clusters was found. Our naming follows the form pdbID_chainID. All network views were generated with Cytoscape (Shannon et al., 2003) and CyToStruct (Nepomnyachiy et al., 2015). B) Representative examples of each group. From left to right: alpha-hemolysin (PDB ID 7ahl), TolC (PDB ID 1ek9), BamA (PDB ID 4k3c), NanC (PDB ID 2wjr), PapC (PDB ID 3fip), Hia (PDB ID 2gr7), LptD (PDB ID 5iva).

### Several groups of OMBBs

Our dataset is composed of 130 structurally characterized OMBB proteins which are <85% similar to each other, including 113 which are less than 50% similar to each other. These include single OMBBs, multi-chain OMBBs, and multi-chain lysins, which are used as a control for describing lack of relationship. Our database contains structures from 35 species of bacteria in 25 species of bacteria.

By filtering the results of the HMM profile alignments of our dataset, we searched for meaningful groupings of β-barrels with sequence similarity indicative of an evolutionary relationship. Our groupings are cutoff by a sequence similarity score also known as an E-value. This E-value is the expected number of false positives in the database with a score at least as good as that of the match (Karlin and Altschul, 1990). We find that the OMBBs represent several unique, conserved domains.

At an E-value <1 the HMM alignments distinguish eight, seemingly unrelated groups of OMBBs (fig. 1) – the adhesins, Fim/Ushers, lysins, assembly proteins, efflux pumps, LptD or the lipid assembly proteins, the injectisomes, and the large group of prototypical barrels. These groups are at best weakly related to the prototypical group (table 1). An interactive version of the network, allowing visual inspection of the underlying sequence and structural similarity, is available online (http://cytostruct.info/rachel/barrels/index.html). Because the majority of the barrel structure of the injectisomes is embedded in the periplasmic space and not in the outer membrane, we exclude these from further consideration even though they are described as OMBBs. Figure 1 also excludes any alignments which are exclusively in a non-barrel portion of the OMBB.

In order to find possible common or unrelated ancestors among groups, we searched for sequence and structural alignments between OMBBs and any other structurally solved proteins (fig. 2). An interactive version of the network, allowing visual inspection of the underlying sequence and structural similarity, is available online (http://cytostruct.info/rachel/barrels/pumps.html) In order to see the widest ranges of possible paths, this assessment was done at E value < 20. We find common relatives among all eight groups of barrels except for the lysins and the efflux pumps.

**Table 1.**
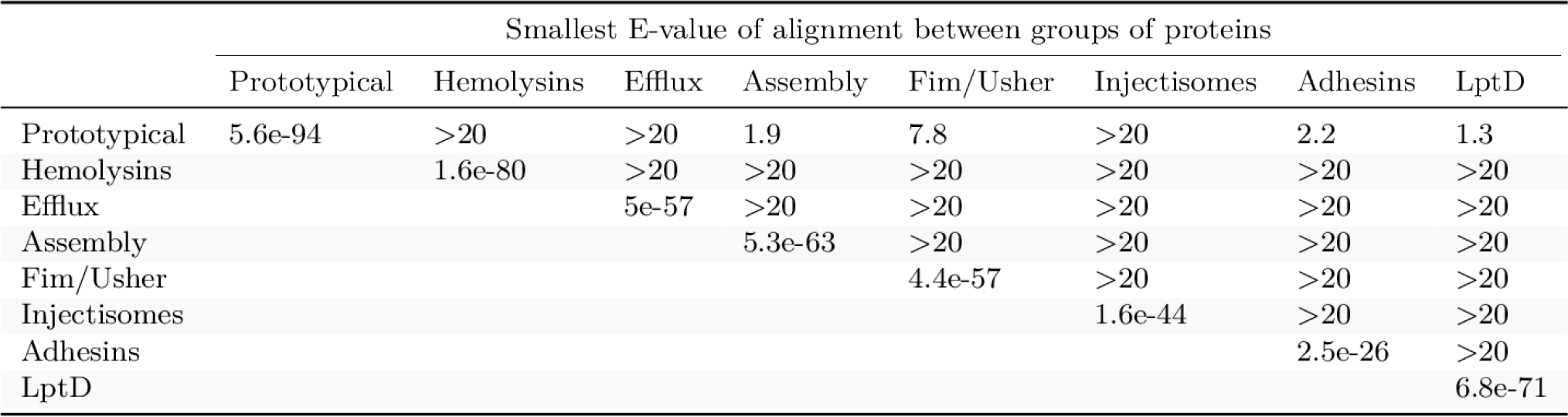
Minimum E-value connecting each group. An E-value of >20 indicates no connection was found.

|  | Smallest E-value of alignment between groups of proteins |  |  |  |  |  |  |  |
| --- | --- | --- | --- | --- | --- | --- | --- | --- |
|  | Prototypical | Hemolysins | Efflux | Assembly | Fim/Usher | Injectisomes | Adhesins | LptD |
| Prototypical | 5.6e-94 | >20 | >20 | 1.9 | 7.8 | >20 | 2.2 | 1.3 |
| Hemolysins |  | 1.6e-80 | >20 | >20 | >20 | >20 | >20 | >20 |
| Efflux |  |  | 5e-57 | >20 | >20 | >20 | >20 | >20 |
| Assembly |  |  |  | 5.3e-63 | >20 | >20 | >20 | >20 |
| Fim/Usher |  |  |  |  | 4.4e-57 | >20 | >20 | >20 |
| Injectisomes |  |  |  |  |  | 1.6e-44 | >20 | >20 |
| Adhesins |  |  |  |  |  |  | 2.5e-26 | >20 |
| LptD |  |  |  |  |  |  |  | 6.8e-71 |

Most of the lysins align with another toxin in the aerolysin family. This is unsurprising given the soluble nature of the monomeric units before barrel formation. Interestingly, the only possible relatives found for the OMBB-containing efflux pumps are the periplasmic adaptor proteins (PAPs) of the same tripartite efflux pumps. Specifically, the helical portions the outer membrane factors (OMFs) of the pumps aligned with the helical portions of the PAPs.

**Figure 2.**
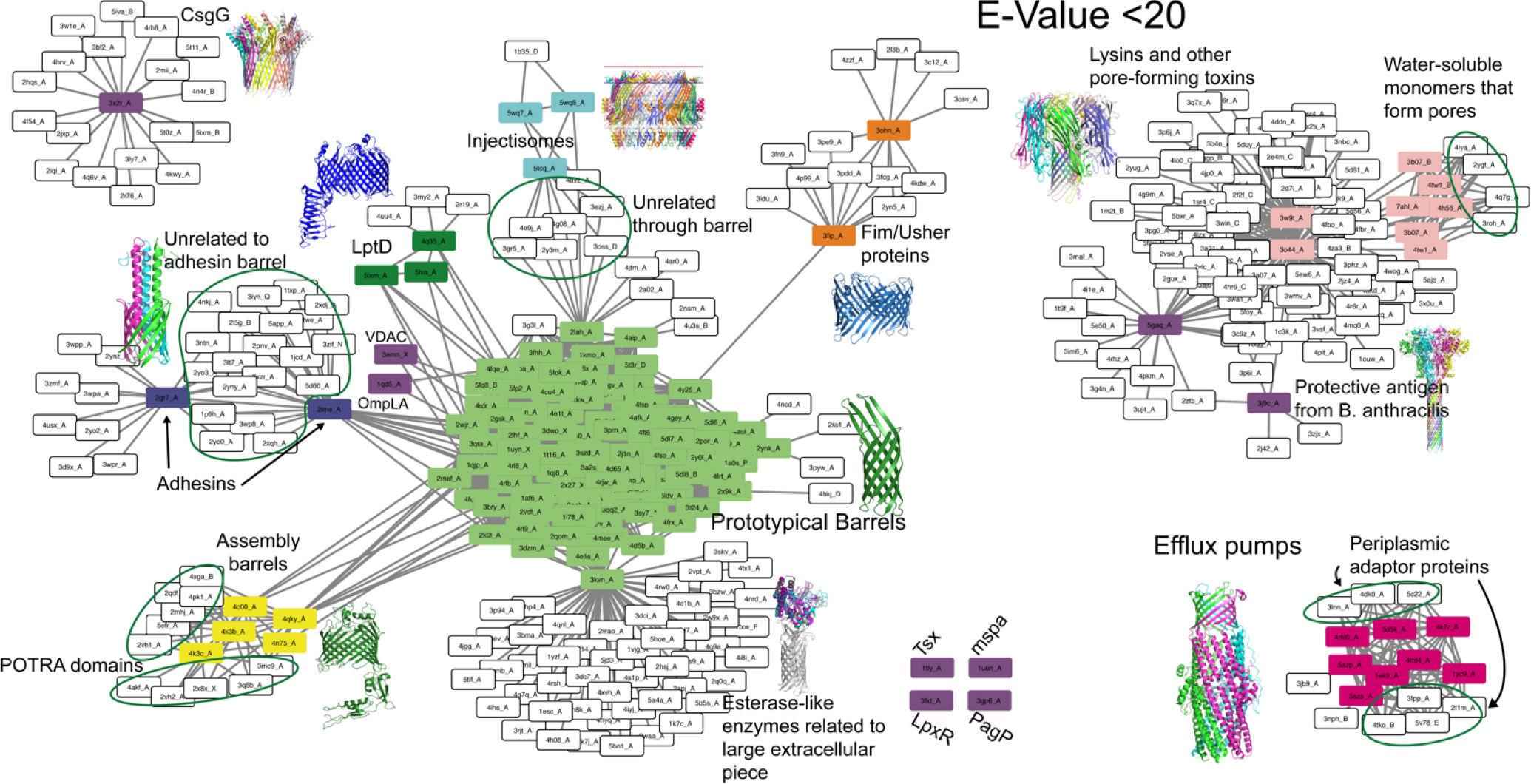
Possible ancestors of OMBB shown through a network of alignments of non-OMBBs to OMBBs at E-value < 20. OMBBs are shown as colored nodes as in Fig 1. White nodes are non-OMBB-containing proteins. Edges represent alignments between two proteins. With a few exceptions primarily involving the lysin cluster or CsgG and discussed in the text, none of the white nodes align with the barrel of an OMBB. The inset structures of the multi-chain OMBBs, such as the efflux pumps and the lysins, are colored by chain.

### Two groups of barrels distinct from the others

Within each of the other six smaller groups (i.e., adhesins, Fim/Usher proteins, lysins, assembly proteins, efflux pumps, and LptDs), the members share a high degree of structural similarity, and have related functions. We find that both the lysin proteins and the efflux pump proteins have more significant structural differences from all other β-barrels.

#### Lysins

Lysins, like alpha-hemolysin and LukGH, are multi-chain barrels with each of either seven or eight chains contributing two strands. The lack of sequence similarity between the lysins and other OMBBs supports the consensus (Remmert et al., 2010) that the lysins are non-homologous to the OMBBs. The lysins are also distinct from the other groups both by organism and by structure. Lysins are the only structures discussed here which are made not only by Gram negative bacteria (De and Olson, 2011), but Gram positive bacteria (Song et al., 1996), as well as by the sea cucumber (Unno et al., 2014). These proteins are exported by their organism of origin and are inserted into the plasma membrane of a target organism (Menestrina et al., 1995). Our structural analysis of these barrels shows recognizable structural differences between these and other barrels. We find that the lysins have a longer barrel than other OMBBs (fig. 3A), with the mean barrel height of 34.2 Å for the prototypical barrels and a mean barrel height of 54.1 Å for the lysins, statistically different from all other barrels with a p-value = 7.23 ×10^−5^ using a Wilcoxon rank-sum test.

#### Efflux pumps

Like the lysins, the efflux pump proteins are multi-chain barrels. However, unlike the lysins, each chain of the efflux pumps contributes four strands. We find this group to be the most structurally different from other barrels. The beta-strands in efflux pumps are disproportionately tilted compared to those of prototypical barrels, at 56.6° compared to 41.7° (fig. 3B and fig. S1) (p-value of 1.46×10^−6^ by Wilcoxon rank-sum test).

**Figure 3.**
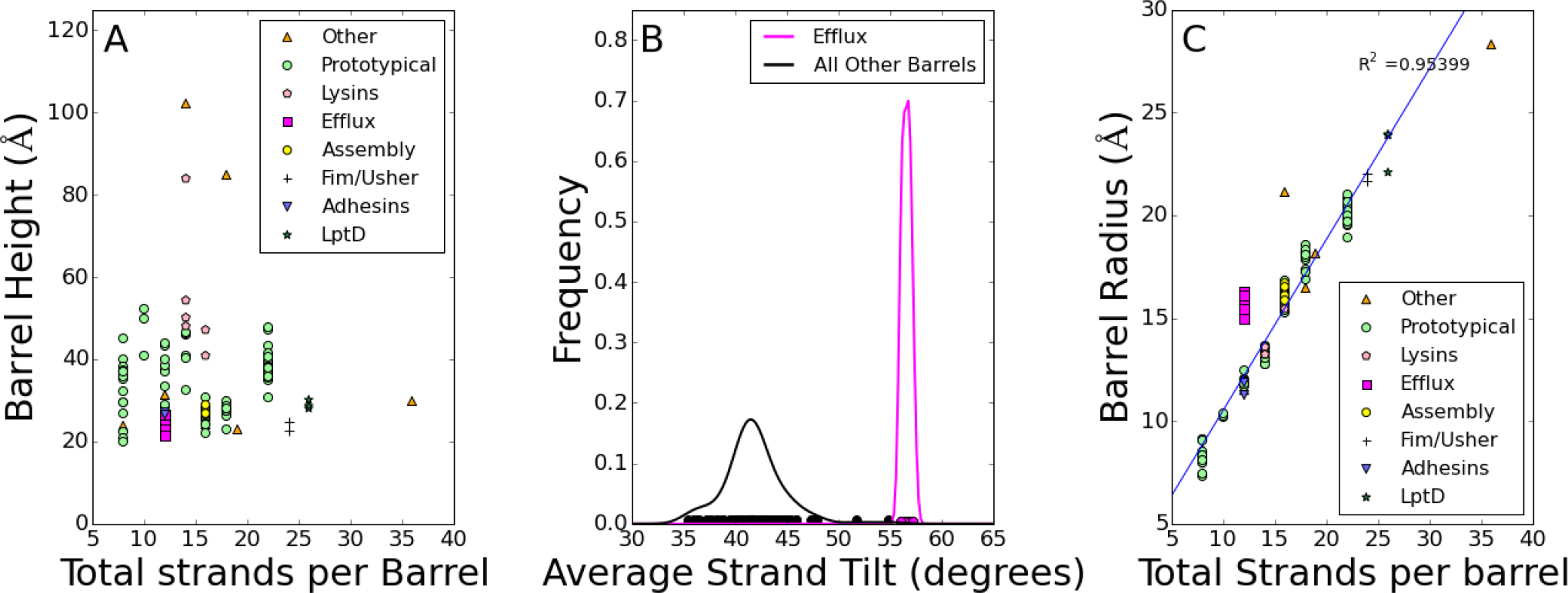
A-C) Structural characteristics of the major components of OMBBs for the six smaller groups of OMBBs, the prototypical OMBBs and the other OMBBs. Groups are colored to match Figure 1. A) Barrel height vs strand number. B) A kernel density estimator was used to show the distribution of each barrel’s average strand tilt in OMBBs. The circles along the bottom represent the data used to generate the kernel. See also fig. S1. C) Graph showing how much each hairpin adds to a barrel’s radius.

In general the addition of each hairpin to a barrel imparts an extra 1.66Å to the radius (fig. 3C). However, as a result of the large tilt, efflux pump OMBBs also have a much wider radius for the total number of strands (n=12) with an average radius of 15.7 Å while the 12-stranded prototypical barrels average 11.9 Å radius. In figure 3C the line of best fit (R^2^ = 0.953) excludes the efflux pump OMBBs. The predicted radius for 12 strands is 12.2 Å which is significantly different from the observed radii in the efflux pump OMBBs with a p-value = 1.235 ×10^−8^ by a paired t-test.

These two characteristics are correlated as the tilt angle dictates the radius. Murzin, Lesk, and Chothia (Murzin et al., 1994) describe a mathematical relationship between tilt and radius, 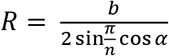, in which b is defined as the interstrand distance (4.4 Å), n is the number of strands, and α is the tilt in radian. We find that this equation holds true for all structures described in all clusters shown in figure 1. Therefore, for a tilt of ~57°, the radius is predicted to be ~15.5 Å. We find no statistical difference between the radius predicted by this formula and the observed values (paired t-test, t = 1.16, p-value = 0.283).

## Discussion

Looking at a combination of structural and sequence information allows us to better understand the different evolutionary paths to OMBBs.

### Convergent evolution of barrels

It is extraordinarily difficult to prove convergence to the exclusion of divergence because bacterial membrane proteins can diverge beyond sequence recognition as described above. Moreover, it has been shown that convergent evolution is not the most likely explanation for many unrelated but structurally similar domains in bacteria (Deeds et al., 2004).

It has previously been hypothesized that the lysins have evolutionarily converged to their barrel structure separately from the divergent evolution of the other β-barrels. This hypothesis was based on the distance in sequence, the difference in structure, and on the differences in organisms that produce them (Remmert et al., 2010). Though other OMBBs in our database are exclusively found in gram negative bacteria, we find one grouping of OMBBs—the efflux pumps— which have no detectable homology to other OMBBs by our metrics.

Not only are the OMBB-containing efflux pumps the least related to the other OMBBs, They also have a subtle but significantly different fold, bolstering our view that it may also have converged on the β-barrel topology. The efflux pumps have more tilted strands and are wider for their chain number than all other barrels studied here. This structural difference along with the lack of sequence similarity leads us to suggest convergent evolution. Moreover, the importance of efflux pumps in allowing bacteria to thrive in hostile environments may have made the evolution of these new types of barrels essential.

The proposed mechanism of an iris-like mode of opening (Andersen et al., 2002) for OMBB-containing efflux pumps may explain both the difference in structure between the efflux pumps and other OMBBs and may also explain the necessity of a separate evolutionary process. Though originally the iris-like mechanism of opening of efflux pumps was believed to only apply to the alpha helical barrel of the OMFs, some have documented movement in the β-barrel as well (Vaccaro et al., 2008). The ability of a β-barrel to modulate its opening for the peristaltic movement of a pump is different from most OMBBs which are understood to be extremely structurally stable and rigid. Evolving a dynamic barrel would be sufficiently different that it could require a separate evolutionary origin.

The alignments between the outer membrane factors of the tripartite efflux pumps and their periplasmic adaptor proteins may be a key to understanding from where the efflux pumps proteins evolved. Efflux pump OMBBs function as the OMF components in tripartate efflux pumps. The OMFs and the PAPs both have barrels of alpha helices that are attached to each other (Fig 4A). It is these regions that have similar sequences to each other. Because the two halves of TolC align with each other (indicating an early duplication event), the two helices of MecA align twice with TolC, once for each half. The helical barrels of MecA and TolC then continue on to beta-sheet regions (Fig 4B). This relationship may be further evidence of the OMBB-containing efflux pumps’ separate origins from the prototypical OMBBs. The idea that the OMFs and the PAPs have evolved from a common ancestor to create a structural palindrome in an efflux pump is appealing. Both would need to be sorted into the periplasm and neither can function without the other. Similar alignments exist between other OMFs and PAPs (Fig 2 and the online network interface http://cytostruct.info/rachel/barrels/pumps.html).

Our theory of having at least two groups of OMBB proteins (efflux pumps and lysins) convergently evolve to such similar folds suggests that there is evolutionary pressure for proteins to find and utilize this fold in the membrane or that this fold is easily ‘designable’ by nature (Koonin et al., 2002).

If the efflux pump β-barrel fold were to be a case of convergent evolution, the question is: what are the pressures that have led to a convergence onto this fold? The membrane barrel topology is fostered by a variety of overlapping factors. The first factor is chemical. As a membrane, the outer membrane precludes long stretches of protein without secondary structure because the hydrophobic membrane cannot satisfy the backbone’s hydrogen bonding requirements. Secondary structure is the primary method of proteins satisfying their own hydrogen bonds. With a defined secondary structure only alpha-helical folds and β-sheet folds are available.

The second factor is biological. Most alpha helical proteins are sorted into the inner membrane. The insertion mechanism of the translocon in the inner membrane causes more uniformly hydrophobic segments to be partitioned into the inner membrane (Osborne et al., 2005). More consistently hydrophobic segments have preference for alpha-helical formation wherein more amino acid side chains have contact with the membrane. Only less hydrophobic segments make it through to the periplasm and allow for folding into the outer membrane. β-barrels fold with two faces, one that can face the membrane and one that can face a proteinaceous or aqueous interior. This favors more hydrophilic membrane proteins. The Bam machinery, especially BamA—itself an OMBB—then assists in insertion of the barrels into the outer membrane through an unknown manner that likely takes advantage of the POTRA domain (Kim et al., 2007) and the weak connection between the first and last strand of BamA (Doerner and Sousa, 2017). Chemically, barrels are preferred over β-sandwiches as barrels allow for complete hydrogen bonding whereas in the β-sandwich architecture the metaphorical crust would have an edge of unsatisfied bonds.

Overall, the sequence and structural relationships among OMBBs demonstrate the possibility that the OMBB fold may have been converged upon from different origins. This convergence on a fold would be facilitated by outer membrane environment requiring β-barrel formation. More work will still need to be done to understand and disentangle the evolutionary pathway of the large group of more clearly inter-related, prototypical OMBBs.

**Figure 4.**
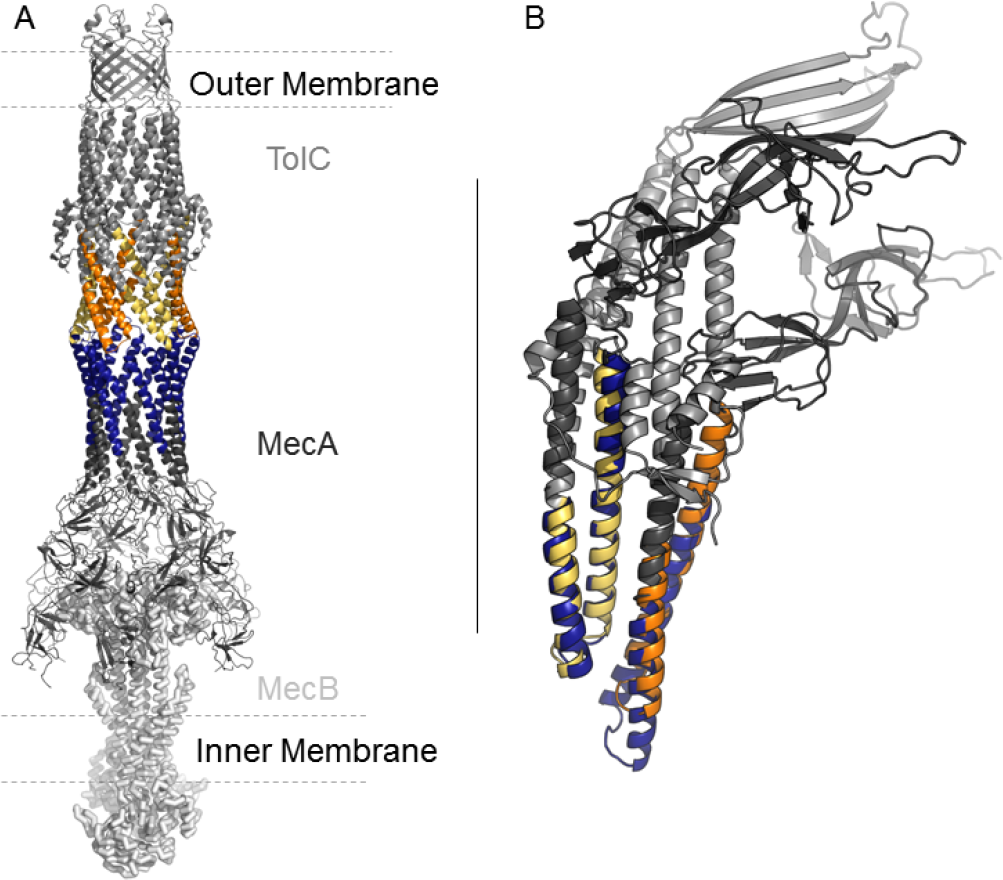
Alignment between TolC and MecA makes a structural palindrome. TolC shown in medium gray, MecA in dark gray, and MecB in light gray. The region where MecA aligns with TolC is shown in blue for MecA and orange and yellow for TolC. Each Each color for TolC represents a different alignment. alignment. **A)** The tripartate MecAB-TolC efflux pump PDB ID 5NIK. **B)** the structural overlay of the two 60 residue sequence alignments between MecA and TolC.

## Acknowledgements

Professors Jamie Walters, Eric Deeds, Christian Ray and Mark Holder for helpful discussions. NIH awards DP2GM128201, P20GM103418, P20GM113117, NSF award TG-MCB160205, The Gordon and Betty Moore Inventor Fellowship, KU-startup and Israel Science Foundation grant number 450/16.

## Author Contributions

Conceptualization, J.S.G.S.; Methodology, J.S.G.S., R.K., M.W.F., and S.N.; Software, M.W.F., S.N., and R.K.; Investigation, M.W.F., R.F., Writing—original draft preparation, J.S.G.S., M.W.F.; Writing-- Review & Editing Preparation, N.B., R.K., J.S.G.S, M.W.F.

## Methods

### Protein Structure Set

We assembled our dataset by compiling native structures from those present in any of the following membrane-focused databases: mpstruc (http://blanco.biomol.uci.edu/mpstruc/, Stephen White lab at UC Irvine), Orientation of Proteins in Membranes or OPM (Lomize et al., 2006), and MemProtMD (Stansfeld et al., 2015). We then used PISCES (Dunbrack and Wang, 2003) to eliminate sequences with greater than 85% sequence identity. This resulted in a set of 110 single-chain β-barrels (including 20 single-chain multi-barrels), and 20 multi-chain barrels: 130 barrels in total. Single-chain barrels are defined as having a single chain contributing to a single complete barrel. In contrast, multi-chain β-barrels have multiple identical chains contributing to a single barrel; only two barrels are heterooligomers (PDB IDs 3b07 and 4tw1) while the rest are homomeric assemblies.

### Strand Definition and Barrel Characteristics

Barrel characteristics were determined using in house software, Polar Bearal, as previously described (Slusky and Dunbrack, 2013). Some updates were included as described below.

Each residue of a chain was classified as belonging to a helix, sheet or other secondary structure using the ϕ and φ angles. Strands were then defined using a combination of backbone hydrogen bonding and a pattern of sheet-like residues. Each barrel was visually checked for appropriate strand definition.

The coordinates of the Cα atoms of the first or last residues of each strand were used to create the top and bottom of the beta-barrel into ellipses (Pauly et al., 2002). The angle between the upper and lower ellipses is the angle between the surface normal of the top and bottom ellipsis. The eigenvectors of the two highest eigenvalues were normalized to form the semi-major and semi-minor of the ellipse, which were then used to calculate the eccentricity. The barrel axis is the vector connecting the centroid of each ellipse while the barrel height is the distance between the two centroids. The radius is the average distance to the barrel axis from the Cα atom of each residue. Barrel tilt is the average angle between the barrel axis and the vector formed between the previous Cα atom and next Cα atom for all residues except the first and last residue in each strand.

### Barrel Network Creation

The barrel network is the relevant sub-network of protein structure space. In it, each node represents a protein, and edges connect proteins that are evolutionary related. We identify evolutionary relationships using the sensitive HMM sequence aligner HHSearch (Soding, 2005), keeping only alignments with a significant score, and sufficient length (Nepomnyachiy et al., 2014, 2017)

HHSearch (Soding, 2005) was used with default parameters to find significant alignments of the set of 130 β-barrels with proteins from the database of 39,386 70% non-redundant PDB chains (April 2017) available with the HHsuite. The profiles were precomputed by the Soding group and downloaded from http://wwwuser.gwdg.de/~compbiol/data/hhsuite/databases/hhsearch_dbs/. HMM profiles for the 57 β-barrels in our list but not in this database were generated using the webserver for HHblits (Remmert et al., 2012) and the database uniclust30_2017_04. The search yielded 2974 alignments at E-value ≤ 1. Alignments with no structural component were removed. We then applied a 20-residue cutoff which is a length that includes 99.7% of the hairpins in our database. All alignments to a non-OMBB structure were visually inspected.

Although it has been shown that using membrane-specific alignment functions yields more accurate alignments for membrane proteins (Stamm et al., 2013), such methods have not yet been applied to β-barrels.

We organized the set of proteins and their alignments as a network, using CyToScape (Saito et al., 2012), and CyToStruct (Nepomnyachiy et al., 2015)). Multiple *alignments* between nodes *frequently* exist, but only the edge with the lowest E-value is shown. To easily view the alignments in a molecular viewer. The resulting network is available online at: http://cytostruct.info/rachel/barrels/. The network including soluble alignments is available online at: http://cytostruct.info/rachel/barrels/pumps.html. In both networks, users can change the E-value cutoff and see the resulting networks: using a more stringent cutoff removes edges, resulting in a more fragmented network. The structure of a protein can be viewed in a molecular viewer by clicking its node. The structural and sequence alignment can be viewed by clicking the edge.

All drawings of proteins were made using Pymol(DeLano, 2002).

